# Loss of *Sun2* ablates nuclear mechanosensing-driven extracellular matrix production and mitigates lung fibrosis

**DOI:** 10.64898/2026.03.18.712778

**Authors:** Emma Carley, Sandra Sandria, Xueyan Peng, Kerri Davidson, Aya Nassereddine, Changwan Ryu, Rachel Rivera, John McGovern, Alexander Ghincea, C. Patrick Lusk, Erica L. Herzog, Valerie Horsley, Megan C. King

## Abstract

Fibrosis and pathological stiffening of tissue are driven by mechanical and biochemical signaling pathways. Here, we find that Sun2, an integral inner nuclear membrane component of Linker of Nucleoskeleton and Cytoskeleton (LINC) complexes, is up-regulated in the lung of patients suffering from fibrotic conditions and in fibroblasts during an injury-induced mouse model of lung fibrosis. Sun2 protein levels also increase in primary lung fibroblasts in a substrate stiffness-dependent manner. *Sun2^-/-^* primary lung fibroblasts respond to TGFβ, become contractile, and express a key marker of extracellular matrix-producing fibroblasts, *Cthrc1*. Consistent with this, Sun2 is dispensable for myofibroblast formation and repairing the alveolar barrier after bleomycin injury. Remarkably, however, fibrosis does not develop in bleomycin-treated *Sun2^-/-^* mouse lungs. This is explained by the requirement for Sun2 to up-regulate genes encoding extracellular matrix proteins. We therefore suggest that Sun2-containing LINC complexes contribute to a mechanical coincidence detection mechanism that acts in concert with canonical TGFβ signaling necessary for pathologic extracellular matrix protein production, representing a nuclear mechanosensing node for intervention in fibrotic diseases of the lung.

## Introduction

Fibrosis is a condition characterized by a pathogenic increase in tissue stiffness that ultimately impairs the function of organs such as the lungs, skin, liver, heart, and kidney (1). Fibrosis is driven by the excessive secretion of extracellular matrix (ECM) proteins by activated fibroblasts (2, 3), often in response to injury (4). No curative therapies currently exist to treat fibrosis, underscoring the need to define the biological mechanisms that drive pathogenesis to inform more effective future treatments (5, 6).

During tissue homeostasis and after injury, signals within the tissue activate quiescent fibroblasts to proliferate, become contractile (transitioning to myofibroblasts) and produce ECM to maintain or rebuild tissue, respectively (7). One of the major drivers of fibrosis across multiple organs is the TGFβ signaling pathway, which acts via Smad transcription factors to regulate genes that orchestrate wound healing including those that encode ECM proteins (4). Recent studies highlight that specific subsets of activated fibroblasts contribute to contractility and collagen production in a manner that reflects their local tissue environment; single cell analyses suggest that contractile, α-smooth muscle actin (αSMA)-positive (*ACTA2*-expressing) fibroblasts and those that maximally produce collagen may be distinct subpopulations (8–13). The persistence or dysregulation of activated fibroblasts can drive a positive feedback loop of tissue stiffening and ECM production to drive fibrotic disease. An ideal therapeutic approach would prevent pathologic collagen production by ECM-secreting fibroblasts while sparing the important contributions of fibroblasts to the wound healing response.

Mechanosensing also plays a key role in driving pro-fibrotic signaling. Cells have multiple mechanisms to sense and respond to changes in their extracellular mechanical environment, including integrin-based focal adhesions, which are mechanosensitive and allow cells to directly sense attributes of the ECM. We showed previously that ECM adhesions engaging β1-integrin drive tension on Linker of Nucleoskeleton and Cytoskeleton (LINC) complexes embedded in the nuclear envelope via the actomyosin cytoskeleton (14). LINC complexes are composed of SUN proteins in the inner nuclear membrane and KASH domain proteins (Nesprins) in the outer nuclear membrane that connect within the nuclear envelope lumen to form a bridge that transmits force across the nuclear envelope (15). Recent evidence suggests that LINC complexes can impact dynamic regulation of fibroblast activation *in vitro* (16).

Here, to determine the mechanisms by which LINC complexes regulate fibrosis, we examined myofibroblast activation, ECM production, and fibrosis development in the lung. We uncover a marked up-regulation of Sun2 protein in fibrotic conditions of the lung in human patients and in mice using the well-defined model of bleomycin-induced lung fibrosis. We demonstrate that substrate mechanics are sufficient to influence Sun2 levels in lung fibroblasts *in vitro*. Ablation of *Sun2* is dispensable for myofibroblast formation, fibroblast contractility, and repair of the alveolar barrier, but it surprisingly blunts pathogenic ECM production *in vivo* and *in vitro*. Thus, Sun2 is required for ECM deposition in response to biophysical cues that are independent of α-smooth muscle actin expression and contractility. Our data implicate Sun2 as a critical regulator linking mechanical stimulation to pathogenic ECM production during fibrosis.

## Results

### The LINC complex component SUN2 is up-regulated in human lung fibrosis

To explore the possible relationship between SUN2 and lung fibrosis, we first analyzed SUN2 levels by immunostaining lung sections from autopsy samples obtained from patients with a common form of fibrotic lung disease, Idiopathic Pulmonary Fibrosis, and from histologically normal lung from patients without known pulmonary disease (Fig. 1A and Supplemental Fig. S1A,B). SUN2 is expressed broadly in the healthy lung parenchyma and the overall level of SUN2 was not different between healthy and IPF tissue (Fig. 1A,B). However, in IPF we observed the highest levels of SUN2 at the nuclear envelope in cells within tissue regions enriched in fibroblasts expressing smooth muscle actin (α-SMA), a marker of contractile, activated fibroblasts that proliferate and reside in dense fibrotic areas (Fig. 1C). Thus, elevated SUN2 could be hallmark of fibrotic lung diseases such as IPF.

**Figure 1.**
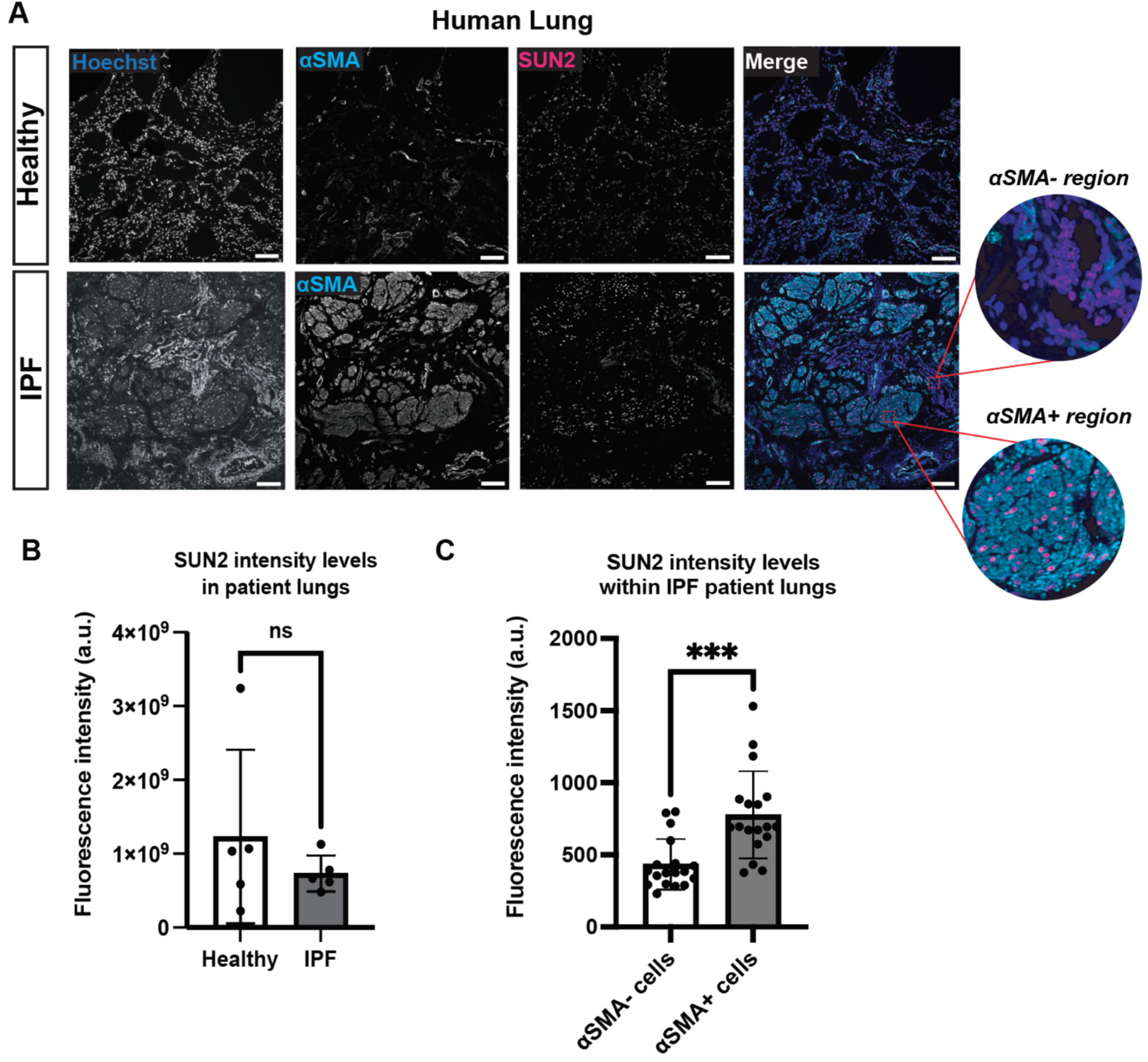
SUN2 is up-regulated in idiopathic lung fibrosis. (**A**) Immunofluorescence staining of alpha-smooth muscle actin (αSMA) and SUN2 in lung tissue samples from healthy human patients and human patients with idiopathic lung fibrosis (IPF). (**B**) SUN2 immunofluorescence staining intensity at the nuclear envelope in healthy and IPF patient lung tissue was quantified in ImageJ, averaging the intensity of >3 fields of view per patient, N=5 patients per condition. No significant difference was found, determined by unpaired two-tailed t test; error bars are SD. (**C**) αSMA+/- cells in IPF patient tissue were identified using QuPath. SUN2 immunofluorescence staining intensity was then measured in both αSMA+/- cells, averaging intensity at the nuclear envelope in cells from >3 fields of view per patient, N=5 patients. There was a significantly higher intensity of SUN2 immunofluorescence staining at the nuclear envelope of αSMA+ cells when compared to αSMA- cells within IPF patient lung tissue. Statistical significance was determined by unpaired two-tailed t test; ***p < 0.001; error bars are SD.

### The nuclear mechanosensing machinery is up-regulated in a murine lung injury model

To determine if the elevated SUN2 levels observed in fibrotic human lungs was recapitulated in a murine model of fibrosis we turned to the well-established bleomycin lung injury model (Fig. 2A). Intratracheal delivery of bleomycin leads to an epithelial cell death response within 48 hours followed by inflammatory injury that transitions within 10 days to fibroblast-driven fibrosis peaking day 14 (17). Comparing the lung parenchyma during injury (Day 2) with the peak of fibrosis (Day 14) we observed a marked up-regulation of Sun2 protein at the nuclear envelope (Fig. 2B,C) at the later timepoint. In alignment with our observations in IPF human lung samples, the levels of Sun2 were higher in alveolar regions positive for α-SMA+ fibroblasts compared to regions enriched in the type I epithelial marker podoplanin (PDPN), indicating that fibroblasts in fibrotic regions express the highest levels of Sun2 (Supplemental Fig. S1C). We similarly observed an increase in the level of Nesprin-1 (Fig. 2D,E), an outer nuclear membrane protein that engages with SUN proteins to constitute LINC complexes. These observations mirror a prior proteomic study demonstrating that both Sun2 and Nesprin-1 levels scale with tissue elasticity across organs (18). Thus, the levels of Sun2 and Nesprin-1, two components of the nuclear mechanosensing machinery, are elevated in response to injury-induced lung fibrosis.

**Figure 2.**
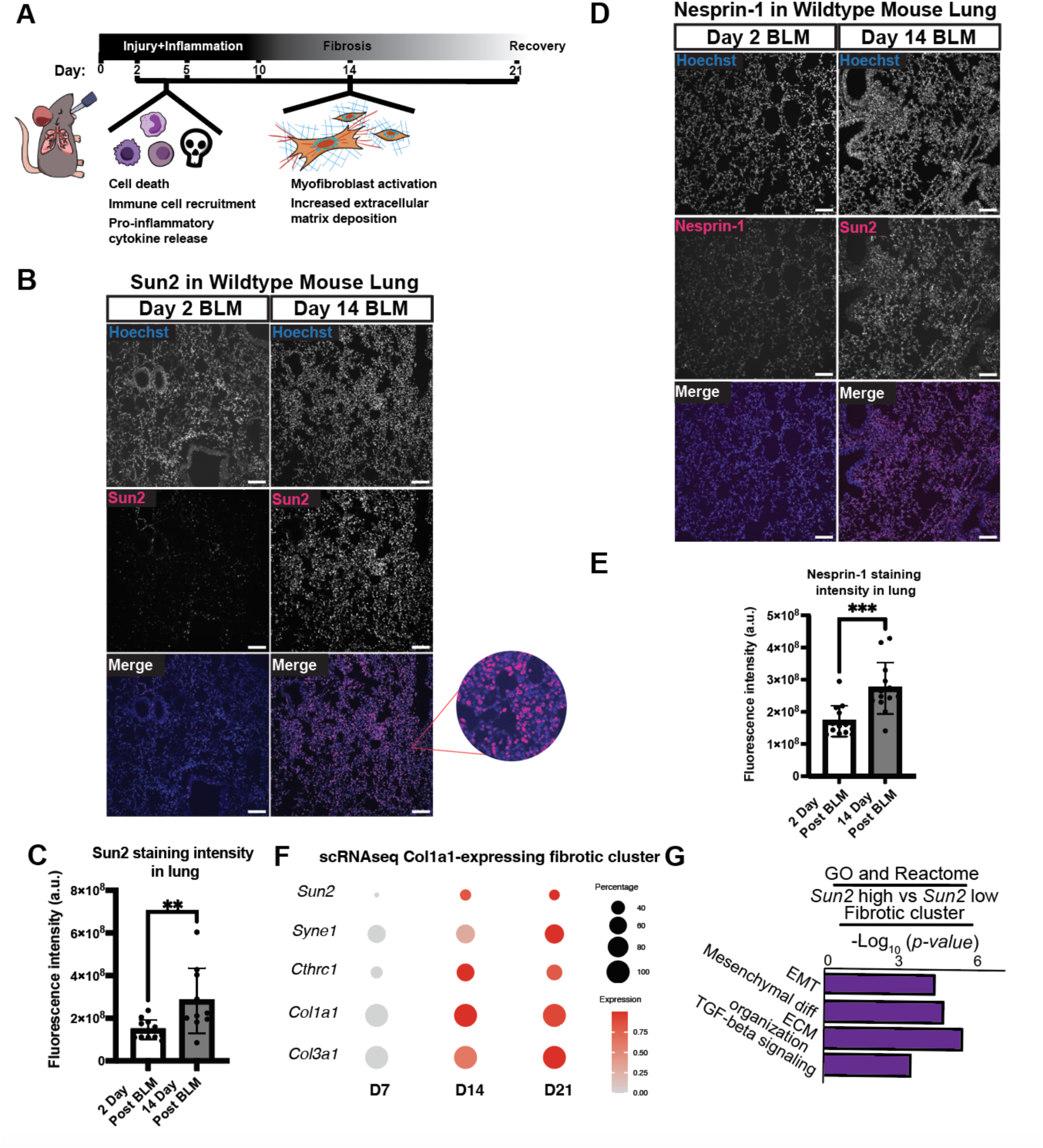
The nuclear mechanosensing machinery is up-regulated in a mouse model of lung fibrosis. (**A**) Schematic demonstrating the time course of bleomycin inhalation induced fibrosis. (**B**) Immunofluorescence staining of Sun2 in wildtype mouse lung tissue 2 days post bleomycin inhalation and 14 days post bleomycin inhalation. (**C**) Sun2 nuclear staining intensity is significantly increased at day 14 post bleomycin inhalation during peak fibrosis when compared to day 2 post bleomycin inhalation. N=3 mice per condition. >2 fields of view analyzed for each mouse. Statistical significance was determined by unpaired two-tailed t test; **p < 0.01; error bars are SD. (**D-E**) Immunofluorescence staining of Nesprin-1 in wildtype mouse lung tissue 2 days and 14 days post bleomycin inhalation. Nesprin-1 nuclear staining intensity is significantly increased at day 14 post bleomycin inhalation during peak fibrosis when compared to day 2 post bleomycin inhalation. N=3 mice per condition, >3 fields of view analyzed for each mouse. Statistical significance was determined by unpaired two-tailed t test; ***p < 0.001; error bars are SD. (**F**) Dot plot of *Sun2*, *Syne1* (encoding Nesprin-1), secretory fibroblast marker *Cthrc1*, and collagens *Col1a1* and *Col3a1* expression in the fibrotic fibroblast cluster of Col1-GFP expressing cells (Supplemental Fig. S2A) from scRNA-seq data in mouse lungs 7, 14, and 21 days after bleomycin inhalation. Data from GEO: GSE210341. (**G**) Gene Ontology (GO) biological process analysis of genes differentially upregulated in cells with high *Sun2* expression compared to those with low *Sun2* expression within the fibrotic cluster.

To explore whether the up-regulation of Sun2 and Nesprin-1 is driven by transcriptional changes we leveraged existing transcriptomic datasets of bleomycin lung injury in mice. Single cell RNA sequencing of Collagen-1-expressing mouse lung fibroblasts in response to bleomycin-induced injury (13) revealed distinct subpopulations, with the highest *Col1a1* expression in the identified “fibrotic” population identified in the UMAP (Supplemental Fig. S2A). In this fibrotic cluster there is an increase in *Sun2* and *Syne1* (encoding Nesprin-1) expression at the peak of fibrosis (Day 14) that persists into the resolution phase (Day 21), concomitant with a marker of highly ECM-secreting fibroblasts (*Cthrc1*) and the collagen genes *Col1a1* and *Col3a1* (Fig. 2F). Leveraging the heterogeneity in *Sun2* expression in fibrotic fibroblasts, we analyzed the differences in gene expression profiles in *Sun2*-high versus *Sun2*-low cells, revealing that higher *Sun2* levels are associated with elevated expression of genes in the GO/Reactome categories of EMT, mesenchymal differentiation, ECM organization and TGFβ signaling (Fig. 2G). Taken together, these data indicate that SUN2 expression by fibroblasts is conserved a fibrotic mechanism in mouse and human lungs.

### Sun2 protein levels are regulated post-transcriptionally in response to substrate stiffness in vitro

To investigate if substrate stiffness is sufficient to increase Sun2 levels, we plated primary mouse lung fibroblasts in soft and stiff conditions *in vitro*. Indeed, we observed higher levels of Sun2 at the nuclear envelope in fibroblasts grown for 24 hours on glass compared to hydrogels of a stiffness that mimics normal lung tissue (2 kPa) (Fig. 3A-B). However, *in vitro* we did not detect an increase in the *Sun2* transcript (Fig. 3C), consistent with prior evidence that Sun2 protein levels are predominantly dictated by post-transcriptional mechanisms (18), including a phospho-degron that regulates Sun2 turnover (19).Thus, fibroblasts exposed to stiff microenvironments activate nuclear mechanosensing machinery.

**Figure 3.**
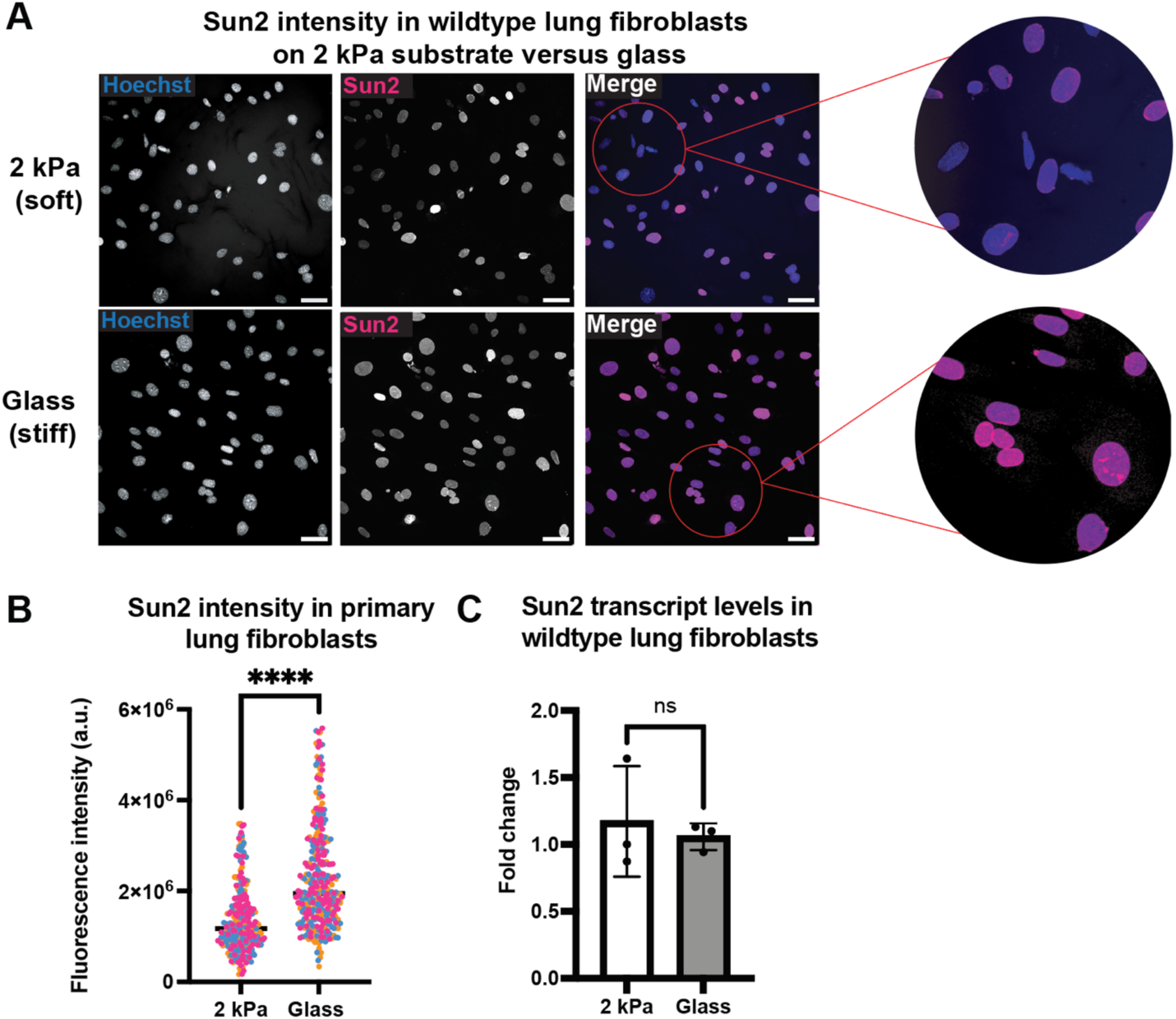
Sun2 protein levels respond to substrate stiffness. (**A**) Immunofluorescence staining of Sun2 in primary wildtype lung fibroblasts seeded on glass and 2 kPa Matrigen substrates for 24 hours. (**B**) Superplot of Sun2 nuclear staining intensity, which is significantly increased in fibroblasts plated on glass compared to a 2 kPa substrate. n=>100 cells, N=3 replicates. Statistical significance was determined by unpaired two-tailed t test; ****p < 0.0001; error bars are SD. (**C**) Plot showing fold change in mRNA expression of *Sun2* in primary wildtype lung fibroblasts cultured on 2 kPa versus glass substrates. There is no significant difference in *Sun2* expression between substrate stiffnesses as determined by two-tailed t test. N=3 replicates. Error bars are SD.

### Sun2 deficiency mitigates bleomycin-induced fibrotic endpoints

To test whether elevated Sun2 levels in fibrotic conditions is required for fibrotic progression, we delivered bleomycin intratracheally to mice with (*Sun2^+/+^)* or without (*Sun2^-/-^*) *Sun2* using a previously described mouse model of constitutive *Sun2* deletion (20). Loss of Sun2 in the lung was confirmed via western blot analysis (Supplementary Fig. S2B). To measure the extent of fibrosis in the *Sun2^-/-^* mice lung, we quantified the levels of soluble, newly-deposited collagen using the Sircol soluble collagen assay of lung lysates after injury (21). While no difference in the amount of soluble collagen was observed at baseline (Day 0) during the inflammatory phase of repair (Days 2 and 5) and early fibrogenesis (Day 10) there were statistically significant decreases in the levels of soluble collagen in the lung tissue at the fibrotic stages of repair (Days 14 and 21) (Fig. 4A). We noticed a bimodal distribution of soluble collagen levels in *Sun2^-/-^* mice at 14-days post-bleomycin injury. Stratifying by sex of the mice revealed that loss of *Sun2* delays the onset of soluble collagen production in males, resulting in significantly less soluble collagen at day 10; in female mice, loss of *Sun2* appears to fully prevent soluble collagen production instead at 14 days (Supplemental Fig. S2C). Thus, there may be a sex difference either in the timing of the manifestation of fibrosis or protection arising from loss of *Sun2,* the relevance of which remains unknown at this time.

**Figure 4.**
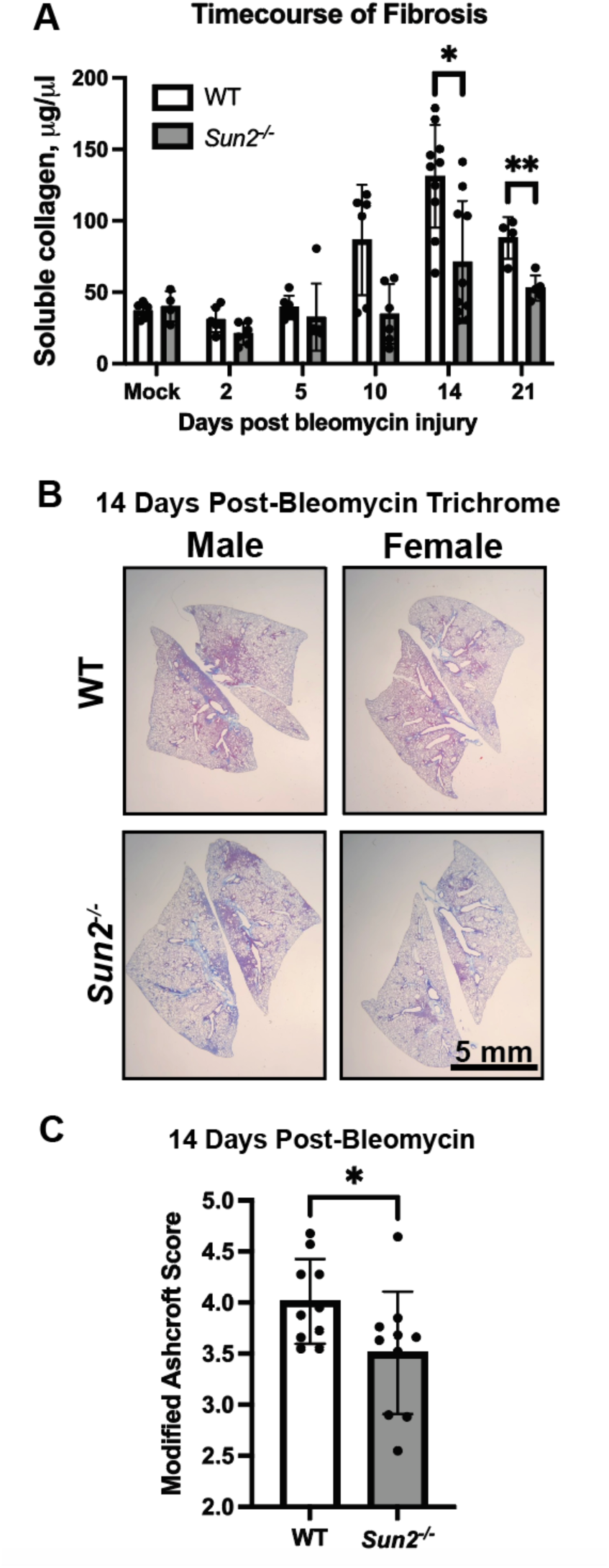
Loss of *Sun2* leads to a decrease in soluble collagen and fibrosis severity after bleomycin-induced lung injury. (**A**) Loss of *Sun2* showed reduced collagen as assessed by Sircol assay at 14 and 21-days post-bleomycin administration. N=>4 mice per condition, male and female mice. Statistical significance was determined by performing multiple t-tests; * p<0.05, ** p<0.01. (**B**) Trichrome staining to visualize collagen (blue) was analyzed to determine disease severity using a (**C**) Modified Ashcroft Score, showing less disease severity in *Sun2^-/-^* mice. N=10 mice per genotype, includes both male and female mice. Statistical significance was determined by performing a two-tailed Student’s *t* test; * p<0.05; Error bars are SD.

We next used Masson’s Trichrome stain to determine if the protective benefit of ablating *Sun2* extends to lung histology (Fig. 4B, Supplemental Figure S2D). Fibrosis severity was evaluated using the semi-quantitative modified Ashcroft scoring (MAS) method described previously (22). Consistent with the Sircol assays results, MAS were reduced in *Sun2^-/-^* lungs 14 days after bleomycin inhalation (Fig. 4C). Together, our data support a paradigm in which loss of *Sun2* protects from severe lung fibrosis..

### Sun2 is dispensable for the initial injury response and re-establishment of the alveolar barrier

It is well established that tissue fibrosis is dependent upon the degree of initial injury (23). To determine whether Sun2 impacts bleomycin-induced cell death and/or the resulting tissue injury we analyzed apoptosis of cells in WT and *Sun2^-/-^* lungs 2 days after bleomycin treatment using TUNEL staining. We observed a non-significant change in TUNEL positive alveolar epithelial cells in mice lacking *Sun2*, indicating that the injury occurred in these mice (Fig. 5A and Supplemental Fig. S3A). To determine if the epithelial barrier was impacted, we analyzed bronchoalveolar lavage (BAL) inflammation and protein content at the day 5 timepoint, which is concomitant with the disruption of the alveolar epithelial barrier (Fig. 5B and Supplemental Fig. S3B). The initial infiltration of immune cells into the BAL was unaffected in *Sun2^-/-^*mice, although there was a modest decrease in the number of cells in the BAL of these animals at 14 days post-injury (Fig. 5B). We further confirmed that reestablishment of the alveolar barrier was normal, and in fact slightly more efficient, in the absence of *Sun2*, by measuring albumin in the BAL (Fig. 5C and Supplemental Fig. S3). These data indicate that *Sun2^-/-^* lung tissue develops bleomycin-induced injury, restores epithelial barrier function, but is protected from a fibrotic response.

**Figure 5.**
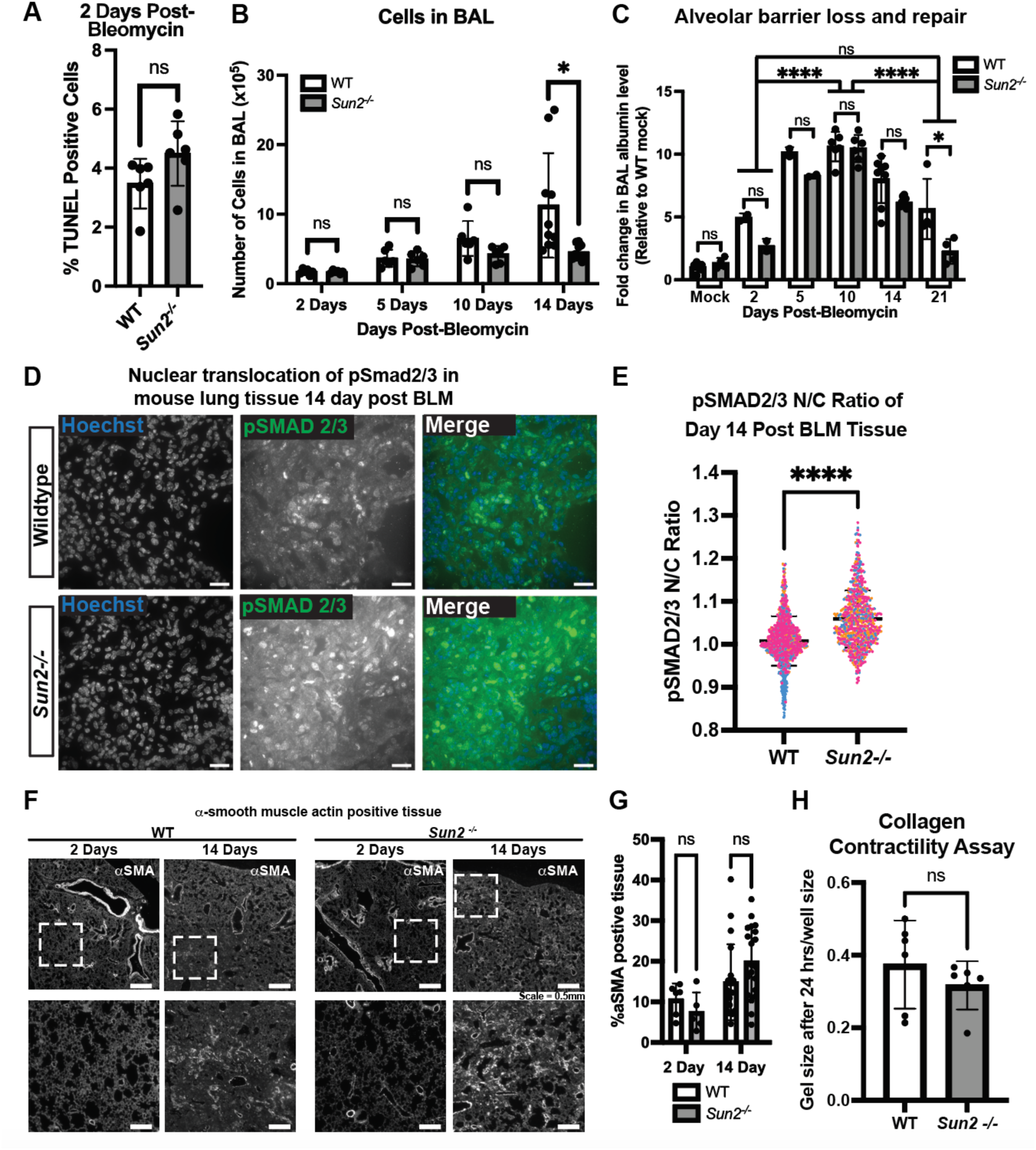
The TGFβ signaling pathway, myofibroblast formation and the ability to heal the alveolar barrier are intact in the absence of Sun2. (**A**) Apoptotic cells were stained using TUNEL and the percentage of apoptotic cells were quantified for tissue 2 days post-bleomycin administration. N=6 male and female mice. There is no statistically significant difference (ns) in these levels upon loss of *Sun2* based on a Student’s *t* test. (**B**) The number of cells present in the BAL is not impacted by the loss of *Sun2* during the inflammatory response phase of the model (prior to Day 14). N=>4 mice per condition, male and female mice. Statistical significance was determined by performing multiple t-tests; ns no significance; * = p<0.05. The Holm–Sidak method was used to correct for multiple comparisons. Error bars are SD. (**C**) Sun2 is dispensable for the recovery of the alveolar barrier as assessed by the kinetics of the increase and subsequent decrease in albumin leaking from the blood into the BAL. (**D**) Wildtype and *Sun2^-/-^* mouse lung tissue from mice euthanized 14 days post bleomycin inhalation were stained with an antibody against phospho-Smad2/3. (**E**) Superplot of the ratio of nuclear to cytoplasmic levels of pSmad2/3 in fibrotic mouse lung tissue shows a significant increase in pSmad2/3 translocation to the nucleus in *Sun2^-/-^* tissue during peak fibrosis. N=3 male mice per genotype, >1 field of view analyzed per mouse. Determined by two-tailed unpaired t test, ****p < 0.0001. Data from three replicates superimposed in blue, orange, and pink. (**F**) Immunofluorescence staining of lung tissue sections from mice 2- or 14-days post-bleomycin administration was performed using an antibody against alpha smooth muscle actin (αSMA) to visualize hypercontractile cells. Regions marked by the dashed box are shown at higher magnification below. (**G**) There is an expected increase in αSMA positive tissue over time but no significant difference in αSMA positive cell levels in WT and *Sun2^-/-^* tissue, as was determined by an ordinary one-way ANOVA followed by Sidak’s multiple comparisons test. Error bars are SD. (**H**) There is no significant difference between the size of the gel contracted by wildtype lung fibroblasts and *Sun2^-/-^* fibroblasts after 24 hours as determined by two-tailed unpaired t-test. N=6 replicates. Error bars are SD.

### Sun2 is dispensable for canonical TGFβ signaling and the fibroblast to myofibroblast transition

To gain more insight into the response to injury we further interrogated TGFβ signaling. Loss of *Sun2* did not impact the production, release, and activation of TGFβ1, as its concentrations in the BAL of mice were unaffected by loss of *Sun2* (Supplemental Fig. S4A). In addition, immunofluorescence staining with antibodies to activated, phospho-Smad2/3 (pSmad2/3) in WT and *Sun2^-/-^* lung tissue demonstrated an increase in the nuclear to cytoplasmic ratio and higher overall pSmad2/3 levels at the peak of fibrosis (Day 14)(Fig. 5D,E and Supplemental Figure S4B,C). Accordingly, in isolated mouse lung fibroblasts loss of *Sun2* did not affect the accumulation of Smad2/3 in the nucleus in response to exogenous TGFβ1 (Supplemental Fig. S4D,E). These data show that activation of TGFβ1 and its canonical signaling pathway are intact in *Sun2^-/-^* mice.

A key TGFβ-dependent fibroblast behavior that contributes to wound healing is the formation of myofibroblasts, which are typically identified by the expression of alpha smooth-muscle actin (αSMA) that is incorporated into stress fibers. In both *Sun^+/+^* and *Sun2^-/-^*animals we observed abundant αSMA-positive cells in the lung parenchyma during the peak fibrotic response (Fig. 5F,G); this finding was further confirmed in primary mouse lung fibroblasts cultured *in vitro* (Supplemental Fig. S4F). Last, we investigated the contractile phenotype of primary lung fibroblasts from WT and *Sun2^-/-^* mice using collagen gel contractility assays. WT or *Sun2^-/-^* primary lung fibroblasts seeded into collagen gels contracted the matrix to an equivalent extent after 24 hours (Fig. 5H and Supplemental Fig. S4G). Thus, *Sun2* is dispensable for fibroblasts to acquire the αSMA expression and contractile properties that define the myofibroblast state.

### Sun2^-/-^ lung fibroblasts fail to produce pathogenic ECM

In addition to driving fibroblast contractility, TGFβ signaling also contributes to ECM production during fibrosis. Moreover, heterogeneity in fibroblast subsets results in one population with the highest level of αSMA expression (contractile myofibroblasts) and another subset that expresses transcripts for ECM proteins such as Collagen-1a1 (secretory fibroblasts)(8–12). We therefore next asked if *Sun2* impacts ECM gene expression and its dependency on TGFβ1. To this end, we plated WT and *Sun2^-/-^* primary lung fibroblasts cultured on collagen-coated 50 kPa hydrogels that were constructed to mimic fibrotic lung tissue. After treatment with vehicle or TGFβ1 for 24 hours, we performed bulk transcriptome analysis (GEO: GSE325284).

We found that ∼3000 genes were altered in vehicle-treated *Sun2^-/-^*fibroblasts compared to vehicle-treated WT cells. ∼2000 genes were differentially regulated with TGFβ1 stimulation. Interestingly, most of the differentially expressed genes were shared between the two conditions (Supplemental Fig. S4H, Supplemental Table 1). Gene ontology analysis of all changes in the TGFβ1-stimulated condition revealed a strong effect on genes tied to collagen synthesis and turnover (6/27 pathways), while “extracellular matrix production” was the sole enriched pathway for genes uniquely affected by loss of *Sun2* in the TGFβ1-stimulated condition (Supplemental Table 2). Focusing on ECM genes, we observed that loss of *Sun2* enhanced expression of *Cthrc1* (collagen triple helix repeat containing 1), a marker that robustly identifies the highest ECM secreting fibroblasts in prior single cell analyses of bleomycin injury (24)(Fig, 6A). Surprisingly, however, in the absence of *Sun2*, ECM genes including *Col3a1*, *Fn1*, *Tnc*, and *Col1a1*, were all lower in *Sun2^-/-^* fibroblasts. Of note, and consistent with an intact TGFβ signaling pathway in the absence of *Sun2*, all of these pro-fibrotic genes respond to exogenous TGFβ in both genotypes, but the set-point is lower in the absence of *Sun2*. Thus, Sun2 does not alter fibrotic fibroblast formation as indicated by high *Cthrc1* expression but is required for the induction of transcripts that are associated with ECM production during the fibrotic response in response to bleomycin.

Thus far, our data suggest that Sun2 is not required for myofibroblast phenotypes such as contractility and expression of α-SMA and *Cthrc1*, but that Sun2 is necessary to induce ECM gene expression in response to stiff substrates to levels achieved by WT fibroblasts. To further interrogate the impact of Sun2 on ECM production, we analyzed the expression of pro-collagen-a1 in *Sun2^-/-^* lung fibroblasts. While over 80% of WT primary mouse lung fibroblasts in culture express pro-collagen-a1 24 hours after plating on glass as assessed by immunostaining (Fig. 6B), we observed a ∼50% decrease in procollagen-expressing primary lung fibroblasts from *Sun2^-/-^* mice (to ∼40%; Fig.6B,C), reinforcing the transcriptome analysis and the conclusion that Sun2 is required for efficient pro-collagen-1a production in lung fibroblasts. Taken together, our findings identify Sun2 as a key component of a nuclear mechanotransduction cascade that is required for the expression of a subset of TGFβ target genes, specifically those that are up-regulated during pathologic ECM remodeling that drives fibrosis (Fig. 7).

**Figure 6.**
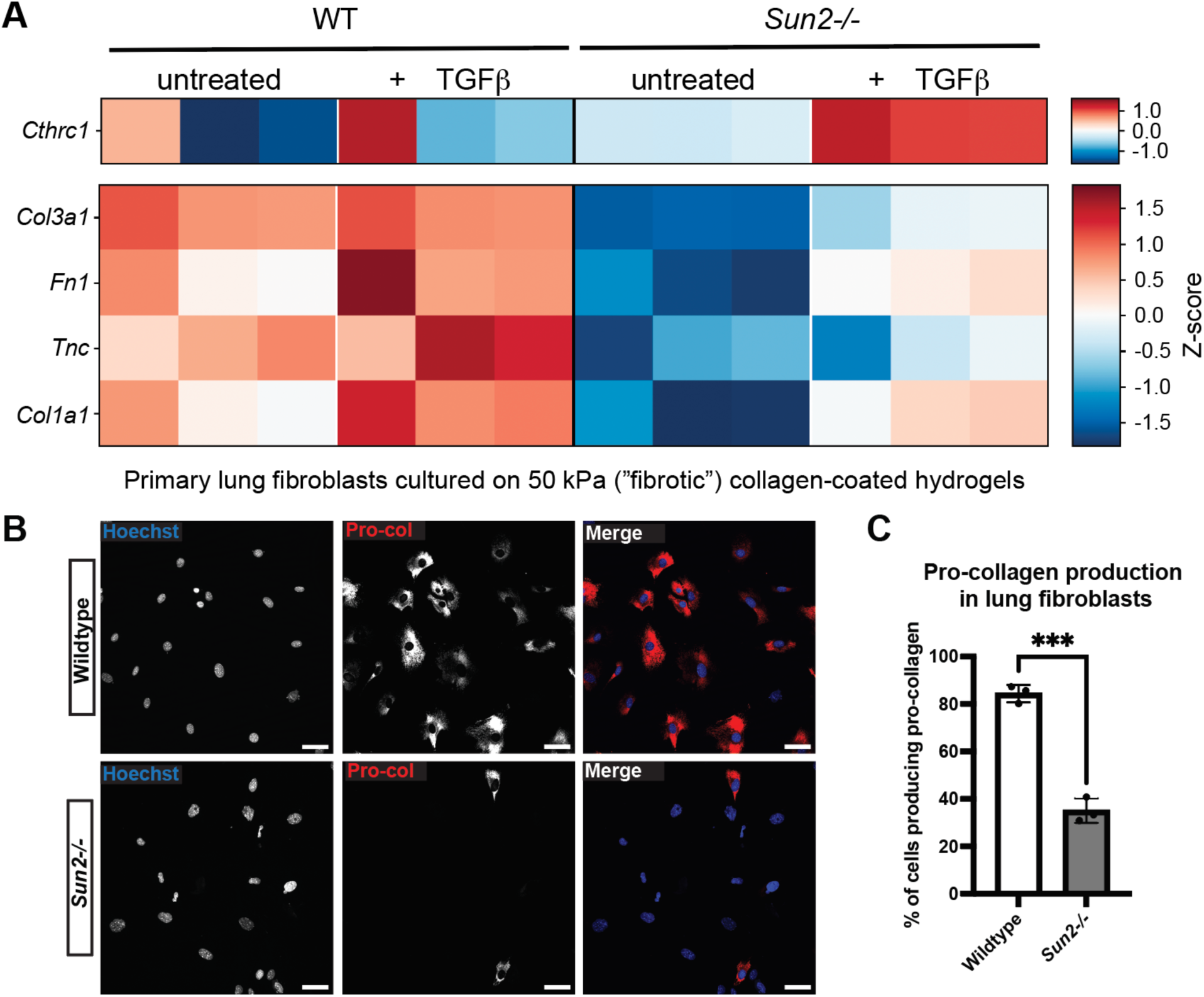
Contractile *Sun2^-/-^*lung fibroblasts fail to express pathogenic ECM. (**A**) Heat map showing expression patterns of genes characteristic of fibrotic fibroblasts in primary wildtype and *Sun2^-/-^* fibroblasts cultured on 50 kPa in the presence and absence of exogenous TGFβ1 (5ng/ml)(GEO: GSE325284). Overall, compared to WT, *Sun2^-/-^* fibroblasts exhibit increased expression of *Cthrc1*, and decreased expression of extracellular matrix genes in the absence or presence of TGFβ1. (**B**) Immunostaining of pro-collagen in primary lung fibroblasts isolated from wildtype and *Sun2^-/-^* mice after growing on glass coverslips for 24 hours. (**C**) Less *Sun2^-/-^* fibroblasts produce pro-collagen. Pro-collagen positivity was determined by the automatic thresholding feature on Fiji. The number of cells positive for pro-collagen staining was counted for each genotype and divided by the total number of cells per replicate and multiplied by 100 to determine a percentage. n=50< cells, N=3 replicates. Determined by two-tailed unpaired t test; ***p < 0.001. (**D**) Cartoon diagram showing our working model in which Sun2 acts as a component of a key nuclear mechanotransduction cascade coupling myofibroblast contractility to collagen production in response to lung injury. In response to injury, subsets of fibroblasts express organized αSMA fibers, contracting surrounding tissue and altering the mechanical environment in both wildtype and *Sun2^-/-^* tissue. Mechanical signals transduced via Sun2-containing LINC complexes in αSMA+ and/or αSMA-fibroblasts prime an increase in transcription of ECM genes. This nuclear mechanosensation is disrupted in *Sun2^-/-^* fibroblasts, which fail to produce pathogenic ECM.

**Figure 7.**
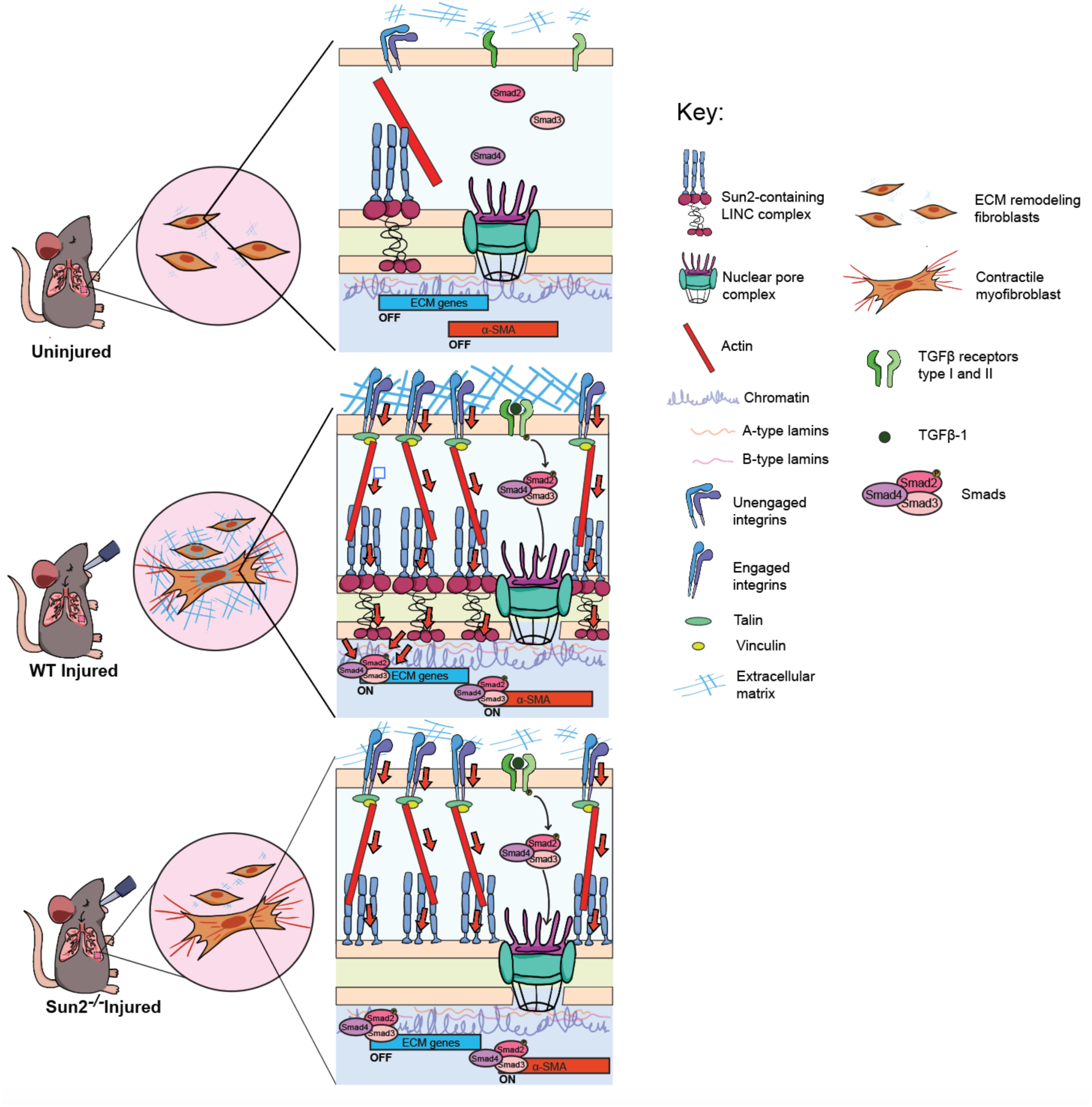
Working model for pro-fibrotic nuclear mechanosensing. Cartoon diagram showing our working model in which Sun2 acts as a component of a key nuclear mechanotransduction cascade coupling a stiffening tissue microenvironment to new collagen deposition in response to lung injury. In the absence of injury, integrins are not strongly engaged, the cytoskeleton is more relaxed, and there is low tension on LINC complexes and the nucleus, corresponding to repression of TGFβ targets including ECM genes and α-SMA. In response to injury, TGFβ binds to the TGFβ receptor and phosphorylated Smads translocate to the nucleus, representing the canonical biochemical cascade. In addition, tension from integrins is propagated through the actin cytoskeleton to LINC complexes. LINC complexes are up-regulated, inducing high tension on the nucleus. Genes contributing to pathogenic ECM production and contractility (ECM genes and α-SMA) are induced, with ECM genes requiring coincidence detection of both nuclear tension and pSmads. The abrogation of nuclear mechanosensation in *Sun2^-/-^* fibroblasts disrupts force propagation into the nucleus, leading to a failure to produce pathogenic ECM but normal contractility (e.g. α-SMA expression) and protection from fibrosis.

## Discussion

In this study, we introduce Sun2 as a key factor that promotes the progression of fibrosis, suggesting that disrupting nuclear mechanotransduction may be an axis for the design of future anti-fibrotic therapies. Our data suggest that Sun2 acts in parallel with TGFβ1, a well-established pro-fibrotic cytokine, to drive fibrosis. This model is supported by four lines of evidence: 1) Bioactive TGFβ1 concentrations in the BAL are similar between WT and *Sun2^-/-^* animals at 10 or 14 days after bleomycin injury (Supplementary Fig. S4A); 2) the expression and nuclear localization of Smads is not reduced but instead elevated in *Sun2^-/-^* bleomycin-treated lung and primary lung fibroblasts (Fig. 5D,E and Supplemental Fig. S4B-E); 3) the transition of fibroblasts to contractile myofibroblasts, a TGFβ1-driven process (4), is Sun2-independent (Fig. 5F-H and Supplemental Fig. S4F,G); and 4) loss of *Sun2* in lung fibroblasts quantitatively decreases expression of ECM genes but retains their responsiveness to TGFβ1 (Fig. 6C). Thus, Sun2 is not required for formation of contractile myofibroblasts, but is necessary to synergize with canonical TGFβ1 signaling to enhance the expression and release of pathogenic ECM by fibrotic fibroblasts.

It is important to note that *Sun2* was not conditionally deleted in this study, and therefore we cannot yet rule out that Sun2 functions in other cell types beyond fibroblasts that impact fibrosis *in vivo*. Indeed, recent work implicates LINC complexes in modulating cell fate determination in the alveolar epithelia. Higher mechanical forces have been reported to promote the AT1 cell fate via chromatin remodeling; in this context full disruption of LINC complexes led to a transition from an AT1 to an AT2 cell state *in vitro* (25). Thus, it remains plausible that some aspect of epithelial repair could be moderately enhanced in *Sun2^-/-^* mice. Ultimately, lineage-specific *Sun2* knockouts and lineage tracing approaches could be informative to fully investigate a possible additional contribution of SUN2 in the epithelia.

Mechanistically, we favor a model in which changes in the mechanics of the lung parenchyma driven by injury-induced myofibroblasts is sensed by Sun2-containing LINC complexes in a distinct, secretory fibroblast subpopulation that are then licensed to produce pathologic levels of ECM (Fig. 6D). This model is consistent with our prior studies showing that 1) *Sun2* uncouples cardiac hypertrophy and fibrosis, which are typically observed concurrently (26); 2) SUN proteins in the epidermis play a key role in transmitting forces from engaged β1-integrin complexes with the ECM to the nucleus to regulate epidermal differentiation (27); and 3) tension on Nesprin-2 is responsive to substrate stiffness (27). The LINC complex is therefore ideally poised to transmit mechanical changes in the ECM to the nucleus. In addition, we find that the Sun2 protein is stabilized in fibrotic conditions (Figs. 1,2), likely in a substrate stiffness-dependent manner (Fig. 3), suggestive of a feed-forward mechanism whereby tissue stiffening will stabilize Sun2-containing LINC complexes, further driving nuclear tension that promotes pathologic levels of new ECM deposition. Such a mechanism could underlie the observation that fibroblasts with disrupted LINC complexes are more plastic and can return to a lower contractile state when moved from a stiff to soft environment more efficiently than control fibroblasts (16). Ultimately, the ability of Sun2 ablation to uncouple productive and pathogenic fibroblast behaviors presents a new opportunity for future anti-fibrotic therapies targeting the nuclear mechanosensing axis.

## Materials and Methods

### Sex as a biological variable

Patients were randomly chosen and were both male and female. Although matched for age and sex between healthy and IPF patients, the limited number of samples prevents conclusions from being drawn on sex as a biological variable. For bleomycin injury experiments we used both male and female animals. While both male and female *Sun2^-/-^* animals showed protection from new collagen deposition, the timing when this reached statistical significance was slightly different as reported in Supplemental Figure S2C. For other experiments including isolation of primary mouse fibroblasts both male and female animals were used; no conclusion can be drawn on sex as a biological variable.

### Human tissues

Human Lung Tissues were provided by The Yale Lung Repository. Tissues provided were archived FFPE specimens that had been previously obtained under Yale’s Rapid Autopsy Protocol from recently deceased persons with no known lung disease or who carried a diagnosis of Idiopathic Pulmonary Fibrosis. These studies are considered non-human subjects research by the Yale IRB.

### Animal handling and tissue isolation

#### Mouse breeding and care

All animal care and experimental procedures were conducted in accordance with the requirements approved by the Institutional Animal Care and Use Committee of Yale University. *Sun2^-/-^* (strain B6;129S6-Sun2tm1Mhan/J) and C57BL/6 WT mice were obtained from Jackson Laboratories. *Sun2^-/-^* mice were previously generated through the replacement of exons 11–16 and part of exon 17 with a neomycin resistance cassette.

#### Bleomycin treatment

Bleomycin administration, bronchoalveolar lavage (BAL), and lung harvest were performed as previously reported (28) in 7-9 week old WT or Sun2-/- mice. Control and treatment groups were maintained under identical conditions. Unless stated otherwise, groups were evenly divided between males and females. Specimens from animals that did not survive to the prespecified endpoints were excluded from analysis.

#### Sacrifice and lung harvest

After animals were terminally anesthetized, they underwent BAL. The lungs were then removed *en bloc* and processed for the requisite assays (21). Whole left lungs were harvested from experimental mice for histological and immunofluorescence analysis. Some samples were embedded in optical cutting temperature (OCT) compound. Others were formalin fixed and paraffin embedded (FFPE).

### Generation of Cell Lines from Mouse Tissue

#### Fibroblast isolation from lung

Fibroblasts were isolated from mouse lung tissue as described (29) with the following modifications. Briefly, WT or Sun2-/- lungs were isolated from 7-9 week-old mice and cut into 1mm pieces. Tissue was digested in 30mL DMEM/F12 media with 1X antibiotic/antimycotic (anti/anti) with 0.13 Wunch units / mL Liberase TM and incubated in a 37C water bath shaker for approximately 30 minutes, until the lung fragments change color from red to white. 20mL DMEM/F12 media with 15% FBS and 1x anti/anti was added to stop the reaction, and the tissue was spun at 500 x g for 5 minutes. The tissue fragments were then washed with 30mL fresh DMEM/F12 media with 15% FBS and 1x anti/anti and spun at 500 x g for 5 minutes twice. Tissue fragments were resuspended in 10mL DMEM/F12 media with 15% FBS and 1x anti/anti and plated in a 10cm dish. Fresh media was added every 7 days until fibroblasts migrated out of the tissue and became confluent. 14 days after isolation, cells were passaged and plated in EMEM with 15% FBS, 1x Penicillin/Streptomycin (Pen/Strep). Cells were incubated in a humidified incubator at 37C, 5% CO2.

### Lung tissue collagen quantification

Lungs were snap frozen in liquid nitrogen and stored at -80C until quantification of collagen by Sircol Collagen Assay according to the manufacturer’s instructions (Zhou et al Sci Transl Med 2014).

### Bronchoalveolar lavage (BAL) analysis

#### BAL cell quantification

BAL cell counts were determined as previously described (Sun et al Arthritis & Rheumatology 2016).

#### BAL Protein Quantification

The concentration of protein in the BAL was quantified using the Pierce BCA protein assay kit (ThermoFisher, 23225) according to the manufacturer’s instructions.

#### BAL SDS-PAGE analysis and albumin quantification

BAL samples were combined 1:1 with 2x SDS-PAGE sample loading buffer and boiled at 95C for 5 minutes. Samples were run in a 4 – 15% SDS-PAGE gel at 200V for 45 minutes. Gels were stained with Coomassie Brilliant Blue overnight and de-stained for 2 hours in 10% acetic acid, 50% methanol in water, then imaged using a scanner. Gray-scale images were analyzed using ImageJ. The integrated density of the bands corresponding to albumin were quantified for each lane and normalized relative to a loading control sample.

#### Cytokine analysis

The levels of TGFβ1 in the mouse BAL was measured using the U-PLEX Custom Biomarker Group 2 Mouse Assays (Mesoscale Discovery) as per manufacturer protocol. Briefly, samples were diluted in a 1:2 ratio and plated in duplicate. Cytokine concentrations were quantified as pg/mL using the Meso Scale Discovery QuickPlex SQ 120 Model 1300 with the Methodical Mind software.

### Protein Sample Preparation and Western Blotting

#### Tissue Protein Samples

5 ug of flash-frozen lung tissue was combined with 500uL ice cold RIPA lysis buffer (150mM NaCl, 1% TritonX100, 0.5% sodium deoxycholate, 0.1% SDS in 50mM Tris HCl pH 8.0, protease inhibitor cocktail 1:100 Sigma P8340), and homogenized on ice using a handheld homogenizer (Kinematica, PT1200E). The lysate was incubated at 4C for 2 hours, with rotation. Lysates were spun at 16000 x g for 20 minutes. The supernatant was combined with 4X sample loading buffer (8% SDS, 40% glycerol, 0.0008% bromophenol blue, 0.4% 2-Mercaptoethanol in 0.25 M Tris HCl, pH6.8) and boiled at 95C for 5 minutes. Protein concentration in the supernatant was measured using the Pierce BCA protein assay kit (ThermoFisher, 23225) according to the manufacturer’s instructions.

#### Cell Culture Protein Samples

Cells were grown to confluence in a 6-well dish. 100uL ice cold RIPA lysis buffer was added to each well, and cells were scraped and transferred to a fresh 1.5mL tube. Cells were incubated on ice for 20 minutes and spun at 4C for 20 minutes. The supernatant was combined with 4X sample loading buffer (8% SDS, 40% glycerol, 0.0008% bromophenol blue, 0.4% 2-Mercaptoethanol in 0.25 M Tris HCl, pH6.8) and boiled at 95C for 5 minutes. Protein concentration in the supernatant was measured using the Pierce BCA protein assay kit (ThermoFisher, 23225) according to the manufacturer’s instructions.

#### Western Blotting

Western blotting was performed as previously described previously (Stewart et al. 2015). Primary antibodies used include SUN2 (Abcam, rabbit, 1:1000, ab123916) and GAPDH (Cell Signaling, rabbit, 1:2000, 2118S).

### Mouse tissue histology

#### Trichrome staining and Modified Ashcroft Score Quantification

Right lung FFPE sections were stained with Masson’s trichrome to visualize collagen deposition. Images of the whole lung were acquired using an iPhone 8 in an iLabCam adaptor attached to a standard light microscope. Images were white balance-adjusted using the Apple “Photos” app version 7.0 for Mac computers. Samples were also imaged using a widefield microscope at 20x magnification for more detailed histological assessment using the modified Ashcroft Score (MAS) (Hübner et al., 2008). Briefly, this quantitative scoring system assigns a numerical value between 0 and 8 that corresponds to the extent of fibrosis in a histologic sample. Multiple regions of each individual sample were imaged and scored, then averaged. Histological analysis of each sample was performed by two individuals blinded to experimental groups.

#### TUNEL staining

Apoptotic cells were detected in paraffin-embedded 5-μm sections from FFPE lung tissue via a TUNEL assay using the ApopTag Peroxidase in Situ Apoptosis Detection Kit (MilliporeSigma, S7100) according to the manufacturer′s protocol, as previously described (Zhou et al., 2014).

### Immunofluorescence

#### Tissue immunofluorescence staining and imaging

Immunofluorescence was performed on both formalin fixed, paraffin embedded (FFPE) sections (5um) and on lung tissue embedded in optimal cutting temperature (OCT) that was cryosectioned (10um). For FFPE sections, the paraffin was removed by incubating the sections in xylene for two 5-minute intervals. Next, sections were re-hydrated by a series of 5-minute incubations in 100% ethanol, 90% ethanol, 75% ethanol, 50% ethanol, and water. Antigen retrieval was then performed by incubating sections in 10mM citrate buffer for 20 minutes in a steamer, then washing samples with PBS. For cryosections, sections were fixed using 4% paraformaldehyde in PBS for 10 minutes, followed by two 5-minute washes in PBS. FFPE and cryosections were treated the same in subsequent steps. Sections were blocked for 1 hour at RT in blocking buffer (3% BSA, 5% donkey serum, 0.05M glycine in PBS2.5% normal goat serum, 1% BSA, 2% gelatin, and 0.25% Triton X-100 in PBS). Sections were incubated with non-mouse primary antibodies in blocking buffer overnight at 4C. The following primary antibodies were used: SUN2 (Abcam; rabbit; 1:1000; ab124916), Nesprin-1 (Abcam; rabbit; 1:100; ab5250), pSMAD 2/3 (Cell Signaling; rabbit: 1:100, D27F4), alpha smooth-muscle actin (Invitrogen; mouse; 1:1000; REF 14-9760-82); pro-surfactant protein C (Abcam; rabbit; 1:1000; ab211326); podopladin (Developmental Studies Hybridoma Bank, hamster, 1:100, 8.1.1-c). Sections were washed with PBS, and a mouse-on-mouse (MOM) kit was used to stain for mouse primary antibodies according to manufacturer’s instructions. Sections were incubated with fluorescent secondary antibodies diluted in PBS 1:2000, and with Hoechst in PBS 1:5000 for 10 minutes. The following secondary antibodies were used: anti-mouse 488 (Invitrogen, A11029); anti-mouse 546 (Invitrogen, A10036); anti-rabbit 647 (Invitrogen, A21245); anti-hamster 488 (Invitrogen, A21110).

Sections were washed with PBS and fluoramount was used to adhere a coverslip. Samples were allowed to dry overnight and were sealed with clear nail polish. Imaging was performed on a Zeiss AxioImager M1 (Zeiss) equipped with an Orca camera (Hamamatsu), a 20x objective, and Zen software (2.6 blue edition; Zeiss). Imaging was performed using the tile scan function and images were stitched using the Zen image analysis program (2.6 blue edition; Zeiss).

#### Fixing and immunostaining of coverslips

Glass coverslips were incubated with 50 ug/ml collagen I from rat tail for one hour at 37 degrees celsius. Collagen I was then washed off with PBS before cells were plated and grown. Experiments done on 2 kPa substrates utilized Matrigen coverslips pre-coated with collagen I. When cells were done growing, they were washed with PBS and fixed with 4% paraformaldehyde in PBS. Cells were permeabilized and blocked with 1% BSA, 0.5% Triton X-100 in PBS for 1 hour at RT. Cells were incubated with primary antibody overnight at 4C. The following primary antibodies were used: alpha smooth-muscle actin (Invitrogen; mouse; 1:1000; REF 14-9760-82); Smad2/3 (Invitrogen; rabbit; 1:1000; 8685S), Pro-Collagen (DSHB; mouse; 1:20; SP1.D8). Cells were washed three times with PBS and incubated with fluorescent secondary antibodies diluted 1:1000 in PBS for 1 hour at room temperature. The following secondary antibodies were used: anti-rabbit 488 (Invitrogen, A11008); anti-mouse 488 (Invitrogen, A11029); anti-mouse 647 (Invitrogen, A31573). Cells were then incubated with Hoechst diluted 1:2000 in PBS, and washed three times in PBS. Coverslips were mounted onto slides using fluoramout, allowed to dry overnight, and sealed with clear nail polish. Imaging was performed on a widefield fluorescence microscope (DeltaVision; Applied Precision/GE healthcare) with a CCD camera (CoolSNAP K4; Photometrics) and SoftWoRx software.

#### Cell line TGFβ1 treatment

2 x 10^5^ WT or *Sun2^-/-^* mouse lung fibroblasts were plated on glass coverslips in a 6-well cell culture plate. The next day, cells were serum starved in their respective media without PBS for 24 hours. Then, cells were treated with 5ng/mL TGFβ1 (Mouse TGF-beta1 Recombinant Protein #15805, Cell Signaling Technology) or an equivalent volume of PBS (mock) in their respective serum-free media for 24 hours before fixing and staining.

### Contractility assay

Adapted from Xie et al. (30), rat collagen I solution (200 µl of 3 mg/ml, Corning® Collagen I, Rat Tail, 100 mg) was mixed with 2.5 × 10^5^ cells in 400 µl DMEM supplemented with 10% FBS (R&D Systems, Minneapolis, MN). 6 ul of 1 M NaOH (6 µl) was added to the cell mixture to initiate gel polymerization. Immediately after NaOH addition, the mixture was homogenized by pipetting up and down and then transferred to a 24-well plate to allow the collagen gel to solidify at 37°C for 15 min. Fresh DMEM (500 µl; 15% FBS) was added to each well and the solidified gel was detached from the well using sterile pipette tips, to allow the gel to float in the culture medium. The gel was incubated at 37 degrees overnight for 24 hours, and then taken out and imaged on a scanner.

### RNA Seq

#### Sample preparation

1 x 10^5^ WT and Sun2^-/-^ mouse lung fibroblasts were seeded in 6 well 50 kPa Matrigen tissue culture plates pre-coated in collagen I from rat tail. Cells were allowed to grow for 24 hours post-seeding in DMEM F/12 15% FBS 1X PenStrep in incubtator at 37° 5% CO2. After these 24 hours, cells were serum starved with DMEM F/12 1% FBS 1X PenStrep for 24 hours. Post serum starvation, cells were then treated with 5 ng/mL TGFβ1 or an equivalent volume of PBS (mock) in their respective serum-free media for 24 hours before RNA isolation.

#### RNA isolation

Total RNA was isolated from WT and Sun2^-/-^ mouse lung fibroblasts using the RNeasy Plus kit (QIAGEN 74034) according to the manufacturer’s instructions for three biological replicates for each experimental condition.

#### Library preparation, sequencing, and analysis

##### RNA Seq Quality Control

RNA quality was determined by estimating the A260/A280 and A260/A230 ratios by NanoDrop (Thermo Fisher Scientific). RNA integrity was determined by running an Agilent Bioanalyzer or Fragment Analyzer gel, which measures the ratio of the ribosomal peaks. Samples with RIN values of 7 or greater were used to prepare libraries.

##### RNA Seq Library Prep

mRNA was purified from a normalized input between 10-1000ng of total RNA using mRNA capture beads prior to fragmentation (Watchmaker mRNA Library Prep Kit). Following first-strand synthesis with random primers, second strand synthesis and A-tailing were performed with dUTP for generating strand-specific sequencing libraries. Adapter ligation with 3’ T overhangs were ligated to library insert fragments. We then selected for amplification of fragments carrying the appropriate adapter sequences at both ends as strands marked with dUTP are not amplified. Indexed libraries were assessed by qRT-PCR using a commercially available kit (KAPA Biosystems) and insert size distribution was determined with the LabChip GX or Agilent TapeStation. Samples with a yield of ≥0.5 ng/μl were sequenced.

##### Flow Cell Preparation and Sequencing

Sample concentrations were normalized to 2.0 nM and loaded onto an Illumina NovaSeq flow cell at a concentration that yields 25 million passing filter clusters per sample. Samples were sequenced using 100bp paired-end sequencing on an Illumina NovaSeq according to Illumina protocols. The 10bp unique dual index was read during additional sequencing reads that automatically follow the completion of read 1. Data generated during sequencing runs were simultaneously transferred to the YCGA high-performance computing cluster. A positive control (prepared bacteriophage Phi X library) provided by Illumina is spiked into every lane at a concentration of 0.3% to monitor sequencing quality in real time.

##### Data Analysis and Storage

Signal intensities were converted to individual base calls during a run using the system’s Real Time Analysis (RTA) software. Base calls were transferred from the machine’s dedicated personal computer to the Yale High Performance Computing cluster via a 1 Gigabit network mount for downstream analysis. Bulk RNA-seq data were then processed using a standard alignment-based workflow. Following initial quality control, adapter sequences were trimmed from raw reads prior to alignment. Reads were aligned to the reference genome using STAR, and gene-level read counts were generated with featureCounts by assigning reads to annotated exons. For downstream analyses, genes were filtered based on total counts across all samples, using a total count cutoff of 100 across all samples for filtering genes. Count-distribution plots were generated before and after filtering to assess the impact of this threshold. Sample-to-sample relationships were evaluated by principal component analysis (PCA) using the first two principal components. Differential expression analysis was performed in R using DESeq2, and results were reported for the Sun2^-/-^ versus WT on 50 kPa comparison and TGFβ treated (as described above) vs untreated conditions.

### RT-qPCR in primary mouse lung fibroblasts

Total RNA was isolated using the RNeasy Plus kit (QIAGEN 74034) according to the manufacturer’s instructions. The iScript cDNA Synthesis Kit (Bio-Rad) was used to generate cDNA from equal amounts of total RNA (1 mg) according to the manufacturer’s instructions. Quantitative real-time PCR was performed with a Bio-Rad CFX96 using iTaq Universal SYBR Green Supermix (Bio-Rad) for 40 cycles. Primers used include GAPDH forward: AAGAAGGTGGTGAAGCAGGC and reverse: TCCACCACCCTGTTGCTGTA; Sun2 forward: ATCCAGACCTTCTATTTCCAGGC and reverse: CCCGGAAGCGGTAGATACAC. Product levels were normalized to GAPDH mRNA levels.

### Image Analysis

#### Quantification of inner nuclear membrane staining intensity

To quantify the intensity of SUN2 and Nesprin-1 immunostaining in human patient and/or mouse lung tissue, image stacks were adjusted to the same brightness and made into sum intensity Z-projections in ImageJ. Hoechst staining was thresholded using the automatic threshold feature of ImageJ. A selection was then created of the threshold and saved as an ROI that was then applied to the SUN2 or Nesprin-1 staining channel of the image. A measurement of integrated density was then taken using the “Measure” feature of ImageJ. Background was accounted for by normalizing to an integrated density measurement of a secondary only control stain of the lung tissue.

#### Quantification of αSMA+ cells in tissue/quantification of SUN2 nuclear intensity in αSMA+ cells

Images of tissue stained with αSMA, or co-stained with αSMA and SUN2, were adjusted to the same brightness in ImageJ and then opened in QuPath (31). Cells were segmented based on Hoechst stain using Stardist (32) and classified as either positive or negative for αSMA using the “create single measurement classifier” object classifier feature in QuPath based on the maximum cytoplasm measurement of the αSMA staining channel. To determine a % of αSMA+ cells in fibrotic tissue four 100 um ROI squares were placed in the lung parenchyma of each field of view of the lung per biological replicate omitting regions with smooth muscle vessels. Nuclear intensity of SUN2 in αSMA+ cells was measured using the add intensity features tool in QuPath.

#### Quantification of Smad2/3 translocation in cell lines

A 5um x 5um region of interest was drawn in ImageJ and used to measure the integrated density of the following: Smad2/3 staining within the nucleus (marked by Hoechst staining), an area of the cytosol proximal to the nucleus, and a measurement of a background, and a cell-free area. The nuclear and cytosolic staining were background-corrected by subtracting the integrated density of the background from that of the nucleus and cytosol measurement. The N:C ratio was calculated by dividing the background corrected values of the nuclear Smad2/3 and the cytosolic Smad2/3 signal.

#### Quantification of pSMAD2/3 nuclear and cytoplasmic intensity and nuclear to cytoplasmic ratio in fibrotic mouse tissue

Images of tissue stained with pSMAD2/3 were adjusted to the same brightness in ImageJ and opened in QuPath. Cells were segmented based on Hoechst stain using Stardist. Intensity measurements in the nucleus and in the cytoplasm were taken using the add intensity feature tool in QuPath. To determine the N/C ratio, nuclear intensity measurements were divided by cytoplasmic intensity measurements.

### Statistics

Statistical tests are described in detail in each figure legend. All indicated statistical tests were performed using GraphPad Prism 9 for macOS (Version 9.4.0).

### Study Approval

All animal care and experimental procedures were conducted in accordance with the requirements approved by the Institutional Animal Care and Use Committee of Yale University.

### Data Availability

Genomic data has been deposited at GEO as GSE325284. All data are publicly available as of the date of publication. This paper also analyzes existing, publicly available data, accessible at GEO: GSE210341. Other data reported in this paper will be shared by the lead contact upon request.

## Author contributions

EC and SS designed, performed and analyzed the majority of the experiments. XP, KD, AN, CR, RR, JM and AG conducted experiments, acquired data, and provided expertise. CPL oversaw project execution and management. ELH, VH and MCK oversaw the study design, execution, management and funding. EC, SS and MCK wrote the manuscript draft, CPL, ELH and VH edited the manuscript, and all authors approved the manuscript prior to submission. The order of the first authors (EC and SS) was alphabetical.

## Acknowledgements

We thank the following individuals for their technical support: Elizabeth Caves, Carrie Ann Davison, Lina Ntokou, and Rebecca Starble. We appreciate the contributions of Dr. Cosmo Saunders, Kayla Hennigan, and Ece Koçak to experiments that contributed to this work but are not included in the manuscript. Many thanks to Elisa Rodriguez for support with mouse husbandry. Yale Pathology Tissue Services prepared samples and performed trichrome staining of FFPE lung. We are grateful to the LusKing lab, Herzog lab, and Nucleus Club at Yale for useful feedback and insights.

## References

1. Rockey DC, Bell PD, and Hill JA. Fibrosis--A Common Pathway to Organ Injury and Failure. N Engl J Med. 2015;373(1):96.

2. Hinz B, Phan SH, Thannickal VJ, Galli A, Bochaton-Piallat ML, and Gabbiani G. The myofibroblast: one function, multiple origins. Am J Pathol. 2007;170(6):1807–16.

3. Pakshir P, Noskovicova N, Lodyga M, Son DO, Schuster R, Goodwin A, et al. The myofibroblast at a glance. J Cell Sci. 2020;133(13).

4. Frangogiannis N. Transforming growth factor-β in tissue fibrosis. J Exp Med.2020;217(3):e20190103.

5. Friedman SL, Sheppard D, Duffield JS, and Violette S. Therapy for fibrotic diseases: nearing the starting line. Sci Transl Med. 2013;5(167):167sr1.

6. Nanthakumar CB, Hatley RJ, Lemma S, Gauldie J, Marshall RP, and Macdonald SJ. Dissecting fibrosis: therapeutic insights from the small-molecule toolbox. Nat Rev Drug Discov. 2015;14(10):693–720.

7. Distler JHW, Györfi AH, Ramanujam M, Whitfield ML, Königshoff M, and Lafyatis R. Shared and distinct mechanisms of fibrosis. Nat Rev Rheumatol. 2019;15(12):705–30.

8. Buechler MB, Pradhan RN, Krishnamurty AT, Cox C, Calviello AK, Wang AW, et al. Cross-tissue organization of the fibroblast lineage. Nature. 2021;593(7860):575–9.

9. Habermann AC, Gutierrez AJ, Bui LT, Yahn SL, Winters NI, Calvi CL, et al. Single-cell RNA sequencing reveals profibrotic roles of distinct epithelial and mesenchymal lineages in pulmonary fibrosis. Science Advances. 2020;6(28):eaba1972.

10. Melms JC, Biermann J, Huang H, Wang Y, Nair A, Tagore S, et al. A molecular single-cell lung atlas of lethal COVID-19. Nature. 2021;595(7865):114–9.

11. Ruiz-Villalba A, Romero JP, Hernández SC, Vilas-Zornoza A, Fortelny N, Castro-Labrador L, et al. Single-Cell RNA Sequencing Analysis Reveals a Crucial Role for CTHRC1 (Collagen Triple Helix Repeat Containing 1) Cardiac Fibroblasts After Myocardial Infarction. Circulation. 2020;142(19):1831–47.

12. Sun KH, Chang Y, Reed NI, and Sheppard D. α-Smooth muscle actin is an inconsistent marker of fibroblasts responsible for force-dependent TGFβ activation or collagen production across multiple models of organ fibrosis. Am J Physiol Lung Cell Mol Physiol. 2016;310(9):L824–36.

13. Tsukui T, Sun KH, Wetter JB, Wilson-Kanamori JR, Hazelwood LA, Henderson NC, et al. Collagen-producing lung cell atlas identifies multiple subsets with distinct localization and relevance to fibrosis. Nat Commun. 2020;11(1):1920.

14. Crisp M, Liu Q, Roux K, Rattner JB, Shanahan C, Burke B, et al. Coupling of the nucleus and cytoplasm: role of the LINC complex. J Cell Biol. 2006;172(1):41–53.

15. King MC. Dynamic regulation of LINC complex composition and function across tissues and contexts. FEBS Lett. 2023;597(22):2823–32.

16. Walker CJ, Crocini C, Ramirez D, Killaars AR, Grim JC, Aguado BA, et al. Nuclear mechanosensing drives chromatin remodelling in persistently activated fibroblasts. Nat Biomed Eng. 2021;5(12):1485–99.

17. Izbicki G, Segel MJ, Christensen TG, Conner MW, and Breuer R. Time course of bleomycin-induced lung fibrosis. Int J Exp Pathol. 2002;83(3):111–9.

18. Swift J, Ivanovska IL, Buxboim A, Harada T, Dingal PC, Pinter J, et al. Nuclear lamin-A scales with tissue stiffness and enhances matrix-directed differentiation. Science. 2013;341(6149):1240104.

19. Krshnan L, Siu WS, Van de Weijer M, Hayward D, Guerrero EN, Gruneberg U, et al. Regulated degradation of the inner nuclear membrane protein SUN2 maintains nuclear envelope architecture and function. Elife. 2022;11.

20. Lei K, Zhang X, Ding X, Guo X, Chen M, Zhu B, et al. SUN1 and SUN2 play critical but partially redundant roles in anchoring nuclei in skeletal muscle cells in mice. Proc Natl Acad Sci U S A. 2009;106(25):10207–12.

21. Zhou Y, Peng H, Sun H, Peng X, Tang C, Gan Y, et al. Chitinase 3-like 1 suppresses injury and promotes fibroproliferative responses in Mammalian lung fibrosis. Sci Transl Med. 2014;6(240):240ra76.

22. !!! INVALID CITATION !!! (22).

23. Lee CG, Cho SJ, Kang MJ, Chapoval SP, Lee PJ, Noble PW, et al. Early growth response gene 1-mediated apoptosis is essential for transforming growth factor beta1-induced pulmonary fibrosis. J Exp Med. 2004;200(3):377–89.

24. !!! INVALID CITATION !!! (13).

25. Shiraishi K, Shah PP, Morley MP, Loebel C, Santini GT, Katzen J, et al. Biophysical forces mediated by respiration maintain lung alveolar epithelial cell fate. Cell. 2023;186(7):1478–92.e15.

26. Stewart RM, Rodriguez EC, and King MC. Ablation of SUN2-containing LINC complexes drives cardiac hypertrophy without interstitial fibrosis. Mol Biol Cell. 2019;30(14):1664–75.

27. Carley E, Stewart RM, Zieman A, Jalilian I, King DE, Zubek A, et al. The LINC complex transmits integrin-dependent tension to the nuclear lamina and represses epidermal differentiation. Elife. 2021;10.

28. Sun H, Zhu Y, Pan H, Chen X, Balestrini JL, Lam TT, et al. Netrin-1 Regulates Fibrocyte Accumulation in the Decellularized Fibrotic Sclerodermatous Lung Microenvironment and in Bleomycin-Induced Pulmonary Fibrosis. Arthritis Rheumatol. 2016;68(5):1251–61.

29. Seluanov A, Vaidya A, and Gorbunova V. Establishing primary adult fibroblast cultures from rodents. J Vis Exp. 2010(44).

30. Xie X, and Percipalle P. Elevated transforming growth factor β signaling activation in β-actin-knockout mouse embryonic fibroblasts enhances myofibroblast features. J Cell Physiol. 2018;233(11):8884–95.

31. Bankhead P, Loughrey MB, Fernández JA, Dombrowski Y, McArt DG, Dunne PD, et al. QuPath: Open source software for digital pathology image analysis. Sci Rep. 2017;7(1):16878.

32. Schmidt U, Weigert M, Broaddus C, and Myers G. Cham: Springer International Publishing; 2018:265–73.

