## Supplemental Figures S1-S4 for "Loss of *Sun2* ablates nuclear mechanosensing-driven extracellular matrix production and mitigates lung fibrosis"

**Carley and Sandria et al., Supplemental Materials**

### Supplemental Figure Legends

**Supplemental Figure 1.** (**A**) Immunostaining of SUN2 and αSMA in lung sections from healthy patients. Three fields of view are displayed for each patient. (**B**) Immunostaining of SUN2 and αSMA in lung sections from IPF patients. Three fields of view are displayed for each patient. (**C**) Immunofluorescence co-staining of Sun2, αSMA, and Pdpn in wildtype mouse lung 14 days post bleomycin shows greater intensity of Sun2 staining in area co-localizing with αSMA.

**Supplemental Figure 2.** (**A**) UMAP plot of scRNA-seq data from Col1-GFP expressing cells in mouse lungs untreated and after 7, 14, and 21 days of bleomycin treatment. Data from GEO: GSE210341. (**B**) Western blot validation of *Sun2*$^{-/-}$ genotype in lung tissue. (**C**) Sircol assay results from Fig. 4A separated by sex at the 10 and 14 days post bleomycin inhalation.

**Supplemental Figure 3.** (**A**) WT and *Sun2-/-* tissue were stained with TUNEL to mark apoptotic cells at 2 days post-bleomycin administration. (**B**) Concentration of protein in the BAL was measured using a BCA assay. As expected, there was a significant increase in levels of protein in the BAL between 2- and 5-days post-bleomycin administration, no statistically significant difference (ns) in protein level upon loss of *Sun2* at any timepoint. Statistical significance was determined by performing an ordinary one-way ANOVA followed by Sidak's multiple comparisons test. ns no significance, *** p<0.0005. Error bars are SD. (**C**) SDS-PAGE followed by Coomassie staining reveal the levels of albumin in BAL samples. As expected, there was a significant increase in levels of albumin in the BAL that peaked at 10 days and began to decrease until levels at 21 days were similar to levels at 2 days post-bleomycin administration. Genotypes of mice, treatment and days after bleomycin inhalation as noted. L= ladder. L=loading control to allow for comparisons between individual SDS-PAGE gels.

**Supplemental Figure 4.** (**A**) Quantification of the total levels of TGFβ1 in the bronchoalveolar lavage (BAL) as measured by ELISA. There is no significant change (ns) in these levels upon loss of *Sun2* at both 10- and 14-days post-bleomycin administration as determined by multiple unpaired t-tests and using the Holm–Sidak method to correct for multiple comparisons. Error bars are SD. (**B**) pSmad2/3 staining is more intense at the nucleus of cells in fibrotic *Sun2-/-* mouse lung tissue. N=3 male mice per genotype, >1 field of view analyzed per mouse. Determined by two-tailed unpaired t test, ****p < 0.0001. Data from three replicates superimposed in blue, orange, and pink. (**C**) pSmad2/3 staining is more intense in the cytoplasm of cells in fibrotic *Sun2-/-* mouse lung tissue. Samples, statistics and plots as in (B). (**D**) Immunofluorescence imaging of Smad2/3 localization (green) in the presence and absence of exogenous TGFβ1 in isolated mouse lung fibroblasts. The nucleus is stained with Hoechst 33342 (blue). (**E**) Quantification of the ratio of nuclear to cytoplasmic levels of Smad2/3 shows no difference in Smad2/3 translocation to the nucleus upon loss of *Sun2*. N=1 replicate, n=>7 cells. Statistical significance was determined by performing multiple t-tests. * = p<0.05, ** = p<0.01. The Holm–Sidak method was used to correct for multiple comparisons. Error bars are SD. (**F**) Immunofluorescence staining of primary lung fibroblasts from wildtype and *Sun2*$^{-/-}$ mice grown on glass coverslips for 24 hours shows establishment of αSMA fibers regardless of genotype. (**G**) To measure contractility of primary lung fibroblasts, cells were embedded in a collagen gel for 24 hours within a 24 well plate. As the fibroblasts contract, the gel (originally the size of the well) shrinks. The gel and well size are outlined for clarity. (**H**) Summary of gene expression changes between cultured WT and *Sun2*$^{-/-}$ mouse lung fibroblasts cultured on 50 kPa hydrogels with and without the addition of TGFβ1. See also Supplemental Tables 1-2.

Supplemental Figure 1.

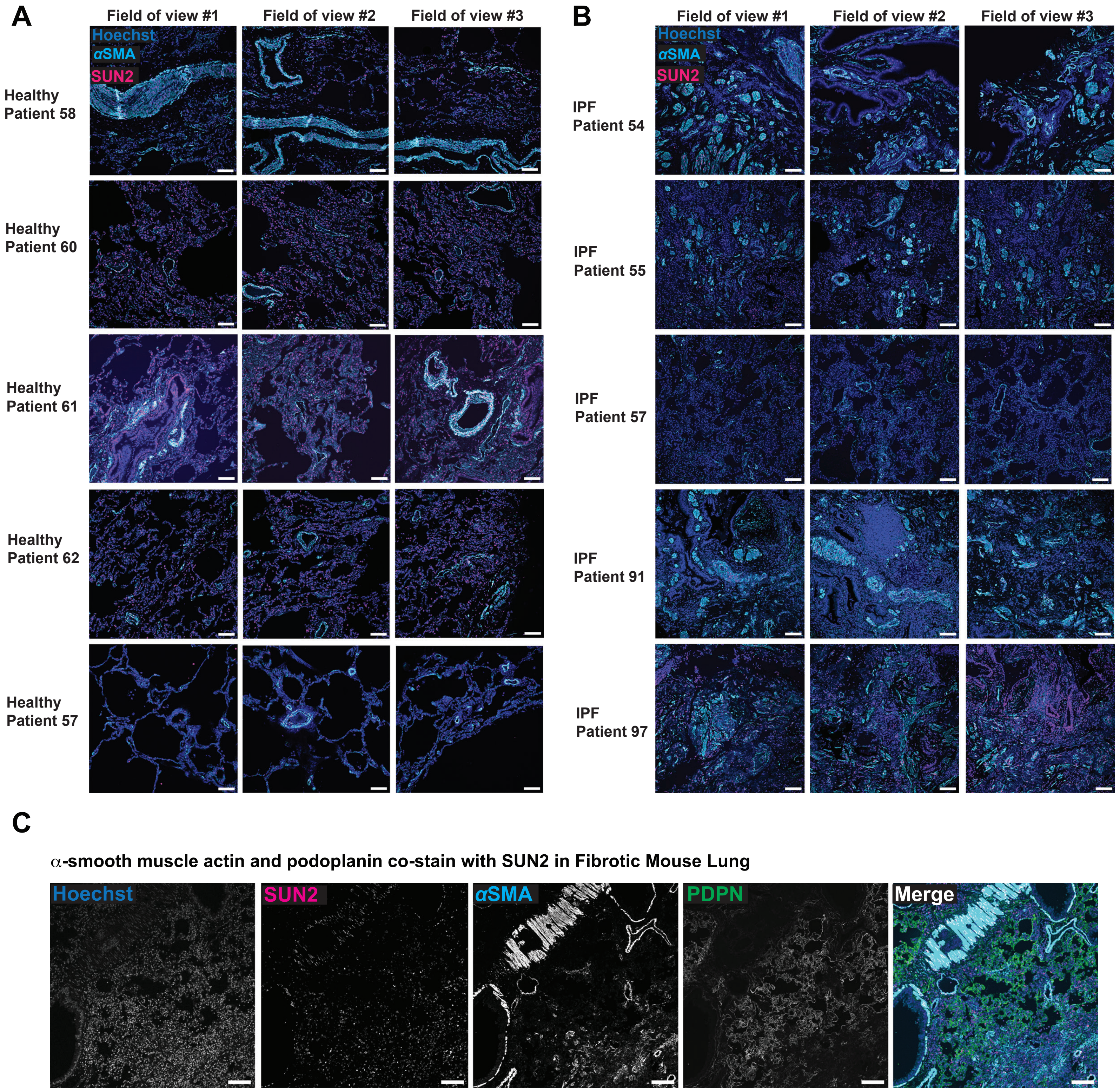

Supplemental Figure 2

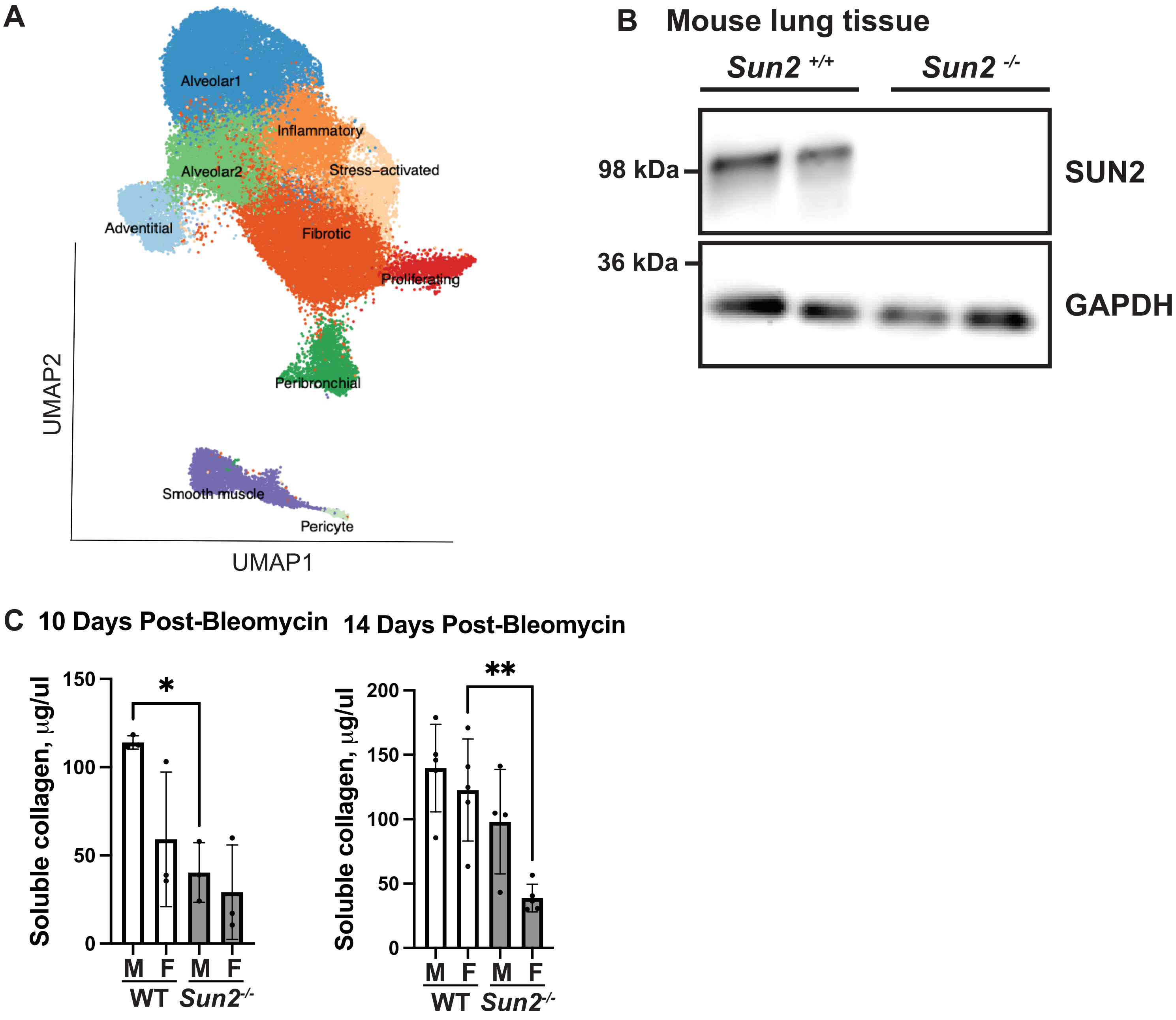

Supplemental Figure 3

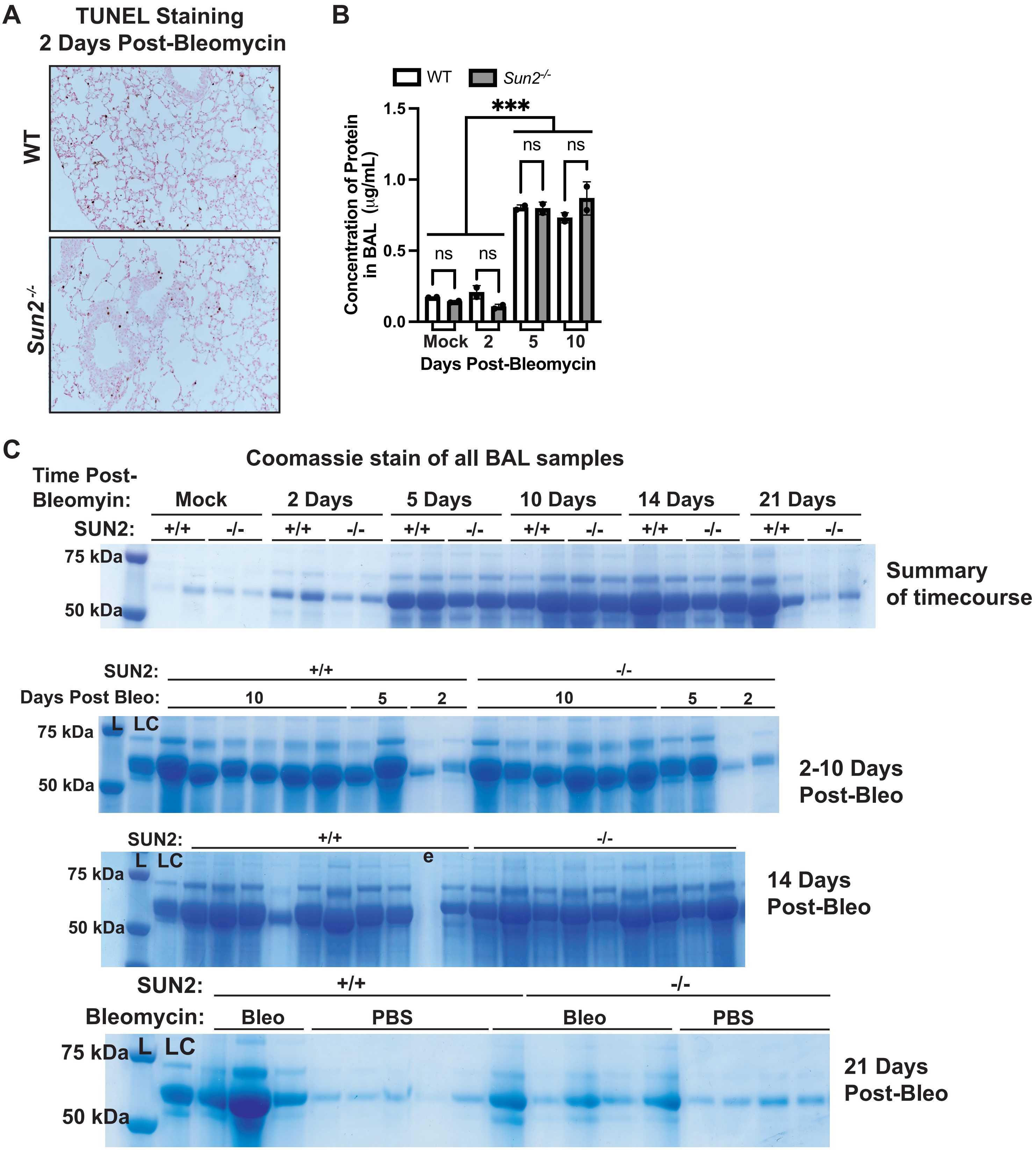

Supplemental Figure S4

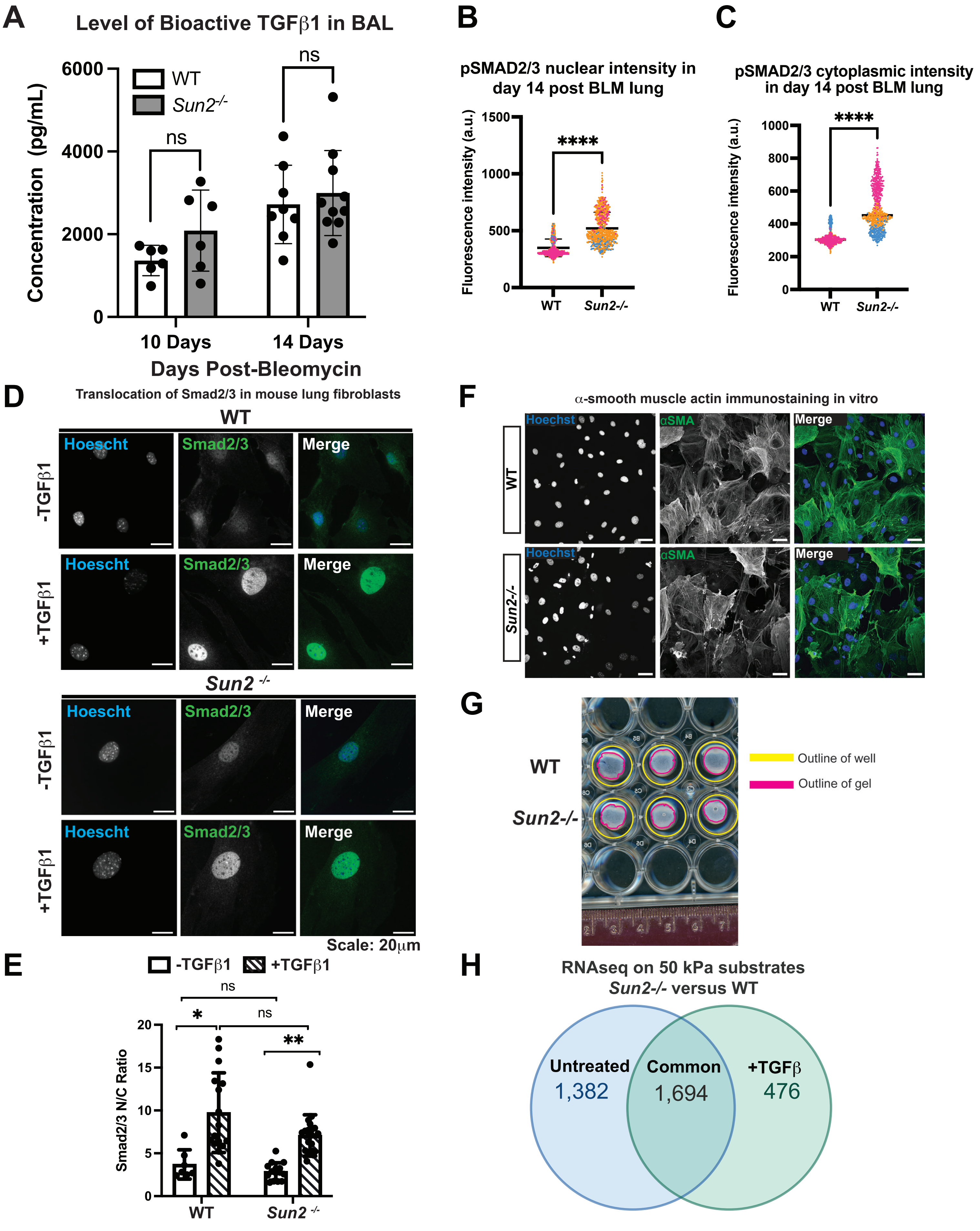
